# Computer Study of Metric Perturbations Impinging on Coupled Axon Tracts

**DOI:** 10.1101/2022.09.20.508763

**Authors:** Aman Chawla, Salvatore Domenic Morgera

## Abstract

Metric perturbations are deviations from a homogeneous spacetime background. In this paper, the authors extend an earlier investigation by using high precision computer simulations and show that there is definite impact of metric perturbations, that is, gravitational radiation, on the time-coded information conducted by a tract of neural axons as found in the human central nervous system.

“… the personal self equals the omnipresent, all-comprehending eternal self … the quintessence of deepest insight into the happenings of the world.”
– E. Schrodinger, *What is Life*?

## 1 Introduction

Gravitational radiation is now detectable via instrumentation on Earth. In contrast to electromagnetic radiation, which faces the barrier of the human skull, gravitational radiation faces no such limitation. In [3], we presented MATLAB simulations showing the impact of gravitational radiation on tracts of human axons, as found in the central nervous system, which had ephaptic coupling between them. We reported there that for strains lower than *h* = 0.09, there was no impact visible on the differential timing of the response of coupled axons. Considering that the gravitational waves received on Earth are about *h* = 1*e* − 21 in strain magnitude or lower [5], we drew the conclusion that there was no impact of gravitational waves on information processing in the brain. In the present article however, we conclude that even weak gravitational waves have a definite impact on information transport by axon tracts.

## 2 Problem Background

The central equation being simulated in this paper is:

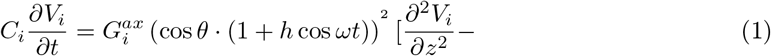

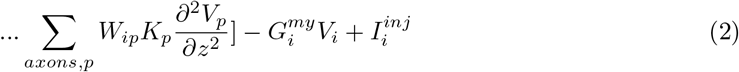

where 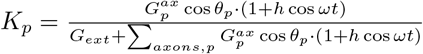 and the notation follows [2].

Given the potential importance of such a first-of-its-kind investigation of gravitational radiation’s impact on a biophysical system (see Figure 1 and Equation (1–2) above), we persisted in seeking some signature of more realistic metric perturbations. We investigated alternate numbers of axons, alternate angular inclinations, alternate inter-axonal distance matrices, along with changes in other parameters including the interfiber resistance to axoplasmic resistance ratio. All of these investigations resulted in the same conclusion – namely, there was no impact at strains below *h* = 0.01. During this period, we didn’t pay attention to the fact that MATLAB’s precision in our implementation was limited to four significant figures which impacted our results. Upon discovering that this precision can be increased to about eighteen significant figures, we were intrigued to re-run our simulations and investigate the output at high precision. As per our expectations at the inception of the project, we do find impact of gravitational radiation on the axon tract action potential initiation time differentials, even at low strains of around *h* = 1*e* − 16.

**Figure 1:**
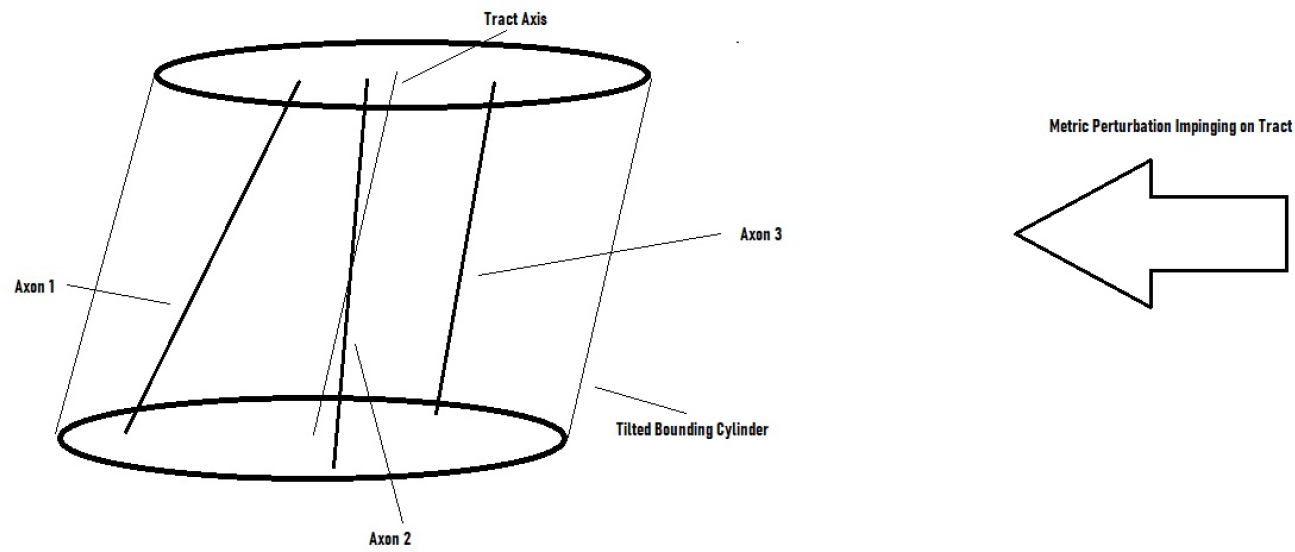
Artistic rendering of a metric perturbation impinging on an axon tract with three axons and an inclined tract axis.

## 3 Results

The medium strain frequency response is shown in Figures 2 through 4. The cosinusoidal term in Equations 1 and 2 has an oscillation-generating impact on the visible traces. The dynamic range of variation of the signals (vertical) present in Figure 2 (angle of 5 degrees) is about 4.5e-8 seconds, in Figure 3 about 7e-8 seconds (angle of 10 degrees) and in Figure 4 about 5e-8 seconds (angle of 15 degrees). This shows a somewhat preferential selection of the 10 degree angle. An analytical approach may illumine why this angle is preferred and it may be related to the orientation of the gravitational wave with respect to this tract’s axons. The medium-low strain frequency response is shown in Figures 5 through 7 and the low strain frequency response is shown in Figures 8 through 10. In each figure, the y-axis is the magnitude of the difference between impulse initiation times (seconds) in axons 2 and 3. The legends indicate the different strain values plotted in colour.

**Figure 2.**
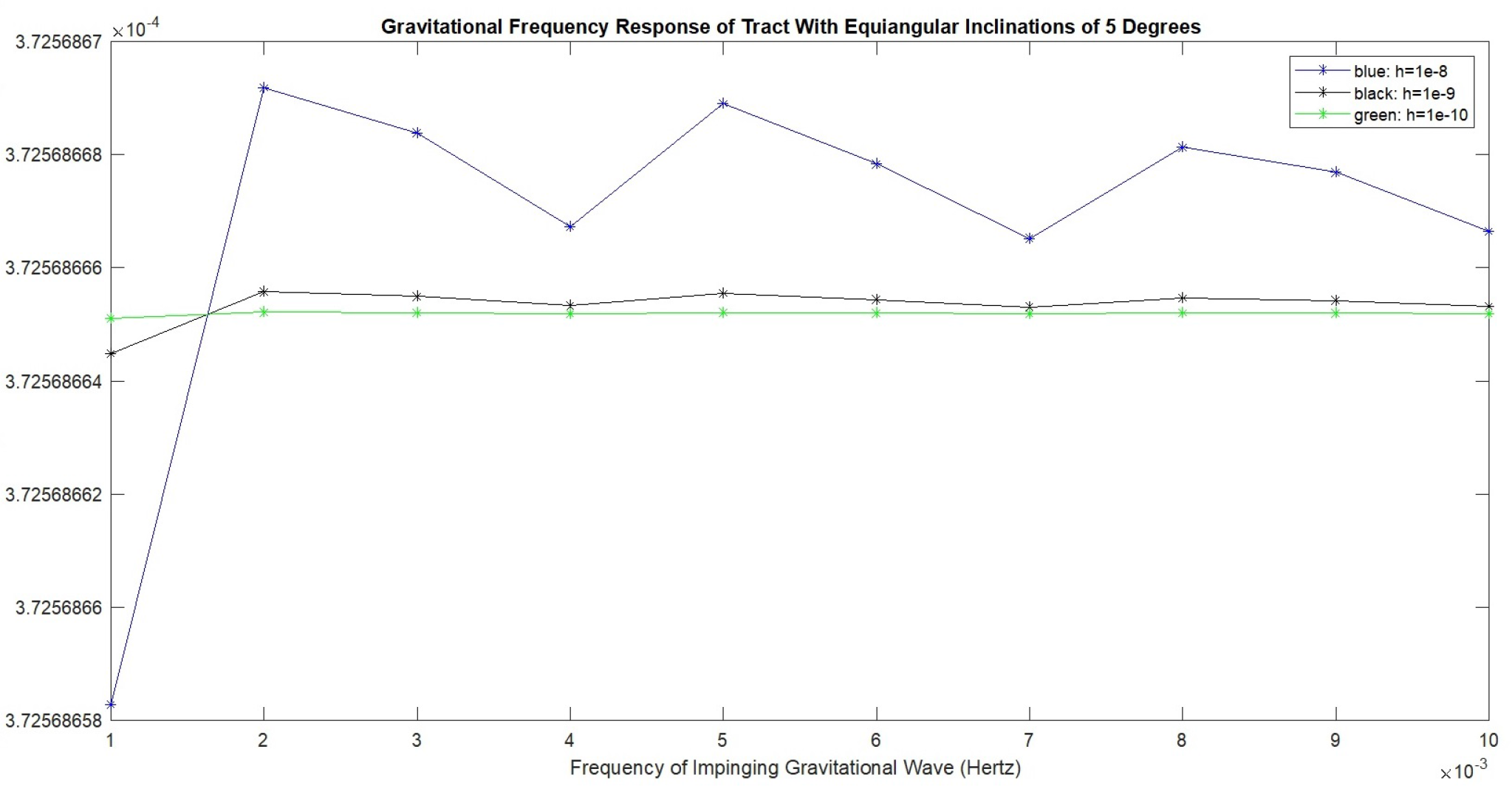
In figure 2, the x-axis is the frequency of the impinging gravitational wave in Hertz, ranging from 1e-3 Hz to 1e-2 Hz. The y-axis is the magnitude of the temporal difference (in seconds) between action potential initiation times on axons 2 and 3 out of a total of 3 axons. Only 1 axon was stimulated, but all 3 are ephaptically coupled within the tract. Each tract considered has a specific geometry. Since the tract was considered in the absence of any ion-channel or other noise source, these are deterministic and not probabilistic simulations. As a result these output graphs are not amenable to statistical significance tests. In each figure, the two applied gravitational wave (dimensionless) strain values are shown in the legend boxes and the point-line graphs are correspondingly color coded. Overall in this set of figures, strains between 1e-8 and 1e-16 are investigated.

**Figure 3.**
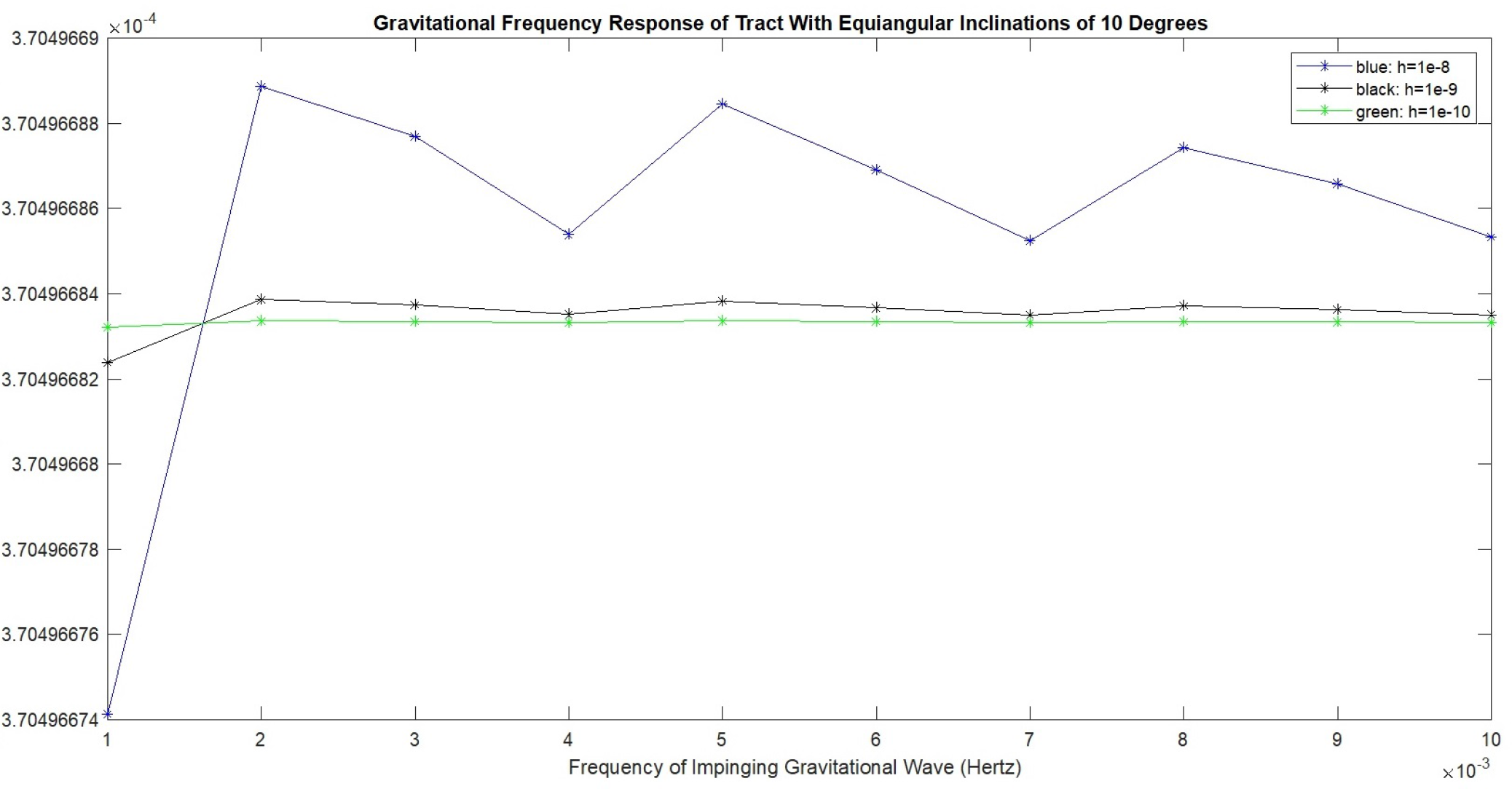
In figure 3, the x-axis is the frequency of the impinging gravitational wave in Hertz, ranging from 1e-3 Hz to 1e-2 Hz. The y-axis is the magnitude of the temporal difference (in seconds) between action potential initiation times on axons 2 and 3 out of a total of 3 axons. Only 1 axon was stimulated, but all 3 are ephaptically coupled within the tract. Each tract considered has a specific geometry. Since the tract was considered in the absence of any ion-channel or other noise source, these are deterministic and not probabilistic simulations. As a result these output graphs are not amenable to statistical significance tests. In each figure, the two applied gravitational wave (dimensionless) strain values are shown in the legend boxes and the point-line graphs are correspondingly color coded. Overall in this set of figures, strains between 1e-8 and 1e-16 are investigated.

**Figure 4.**
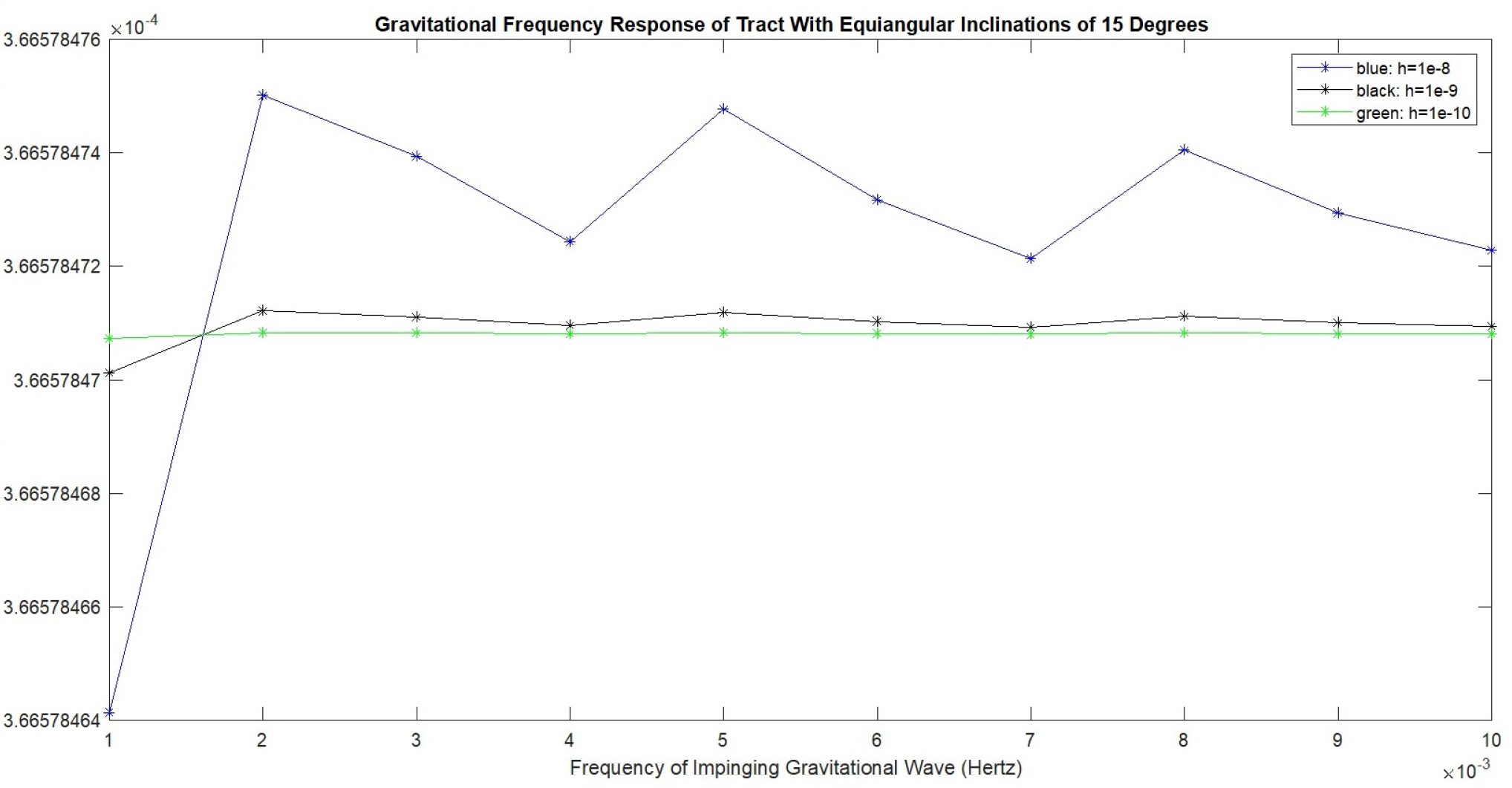
In figure 4, the x-axis is the frequency of the impinging gravitational wave in Hertz, ranging from 1e-3 Hz to 1e-2 Hz. The y-axis is the magnitude of the temporal difference (in seconds) between action potential initiation times on axons 2 and 3 out of a total of 3 axons. Only 1 axon was stimulated, but all 3 are ephaptically coupled within the tract. Each tract considered has a specific geometry. Since the tract was considered in the absence of any ion-channel or other noise source, these are deterministic and not probabilistic simulations. As a result these output graphs are not amenable to statistical significance tests. In each figure, the two applied gravitational wave (dimensionless) strain values are shown in the legend boxes and the point-line graphs are correspondingly color coded. Overall in this set of figures, strains between 1e-8 and 1e-16 are investigated.

**Figure 5.**
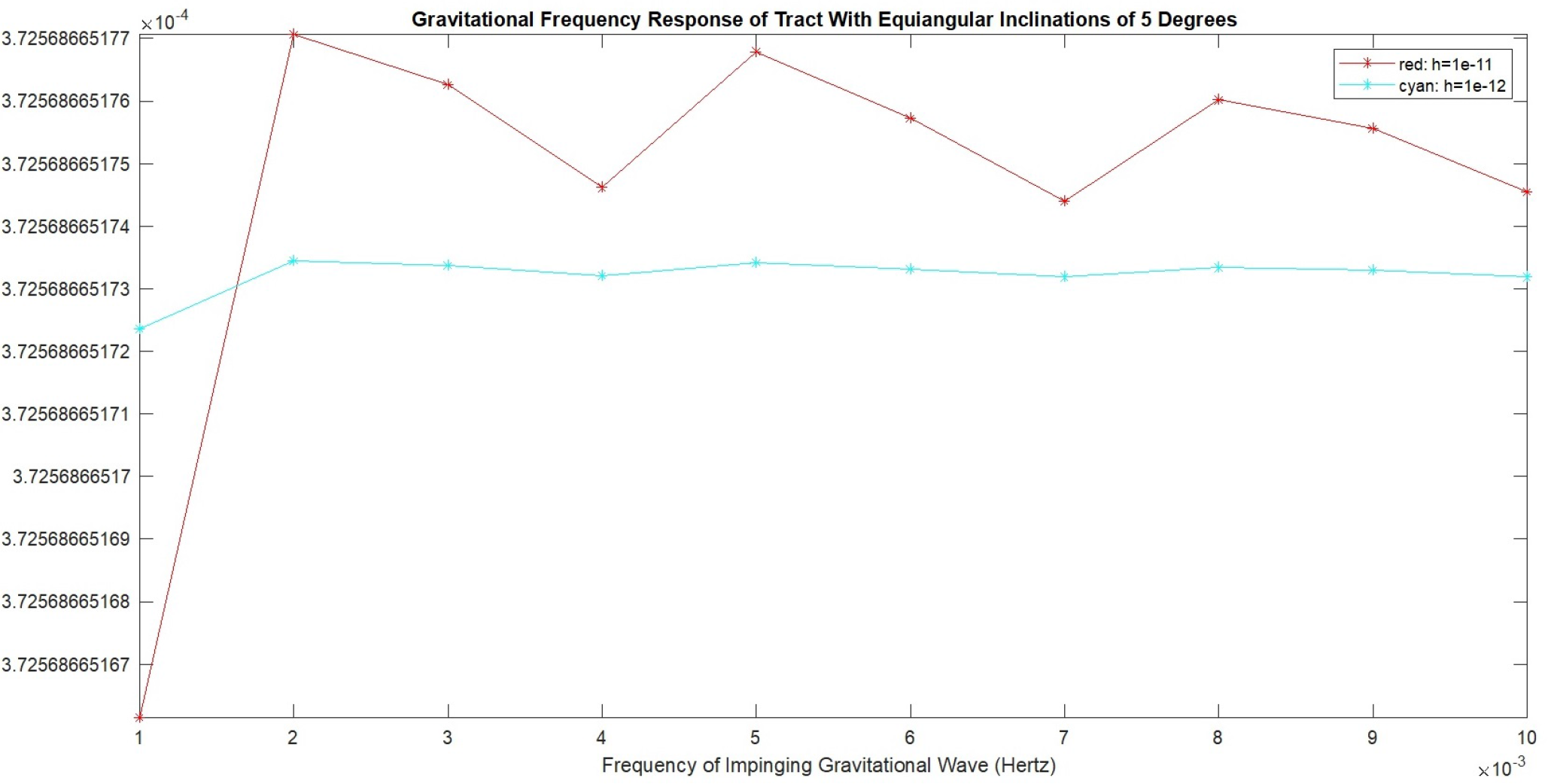
In figure 5, the x-axis is the frequency of the impinging gravitational wave in Hertz, ranging from 1e-3 Hz to 1e-2 Hz. The y-axis is the magnitude of the temporal difference (in seconds) between action potential initiation times on axons 2 and 3 out of a total of 3 axons. Only 1 axon was stimulated, but all 3 are ephaptically coupled within the tract. Each tract considered has a specific geometry. Since the tract was considered in the absence of any ion-channel or other noise source, these are deterministic and not probabilistic simulations. As a result these output graphs are not amenable to statistical significance tests. In each figure, the two applied gravitational wave (dimensionless) strain values are shown in the legend boxes and the point-line graphs are correspondingly color coded. Overall in this set of figures, strains between 1e-8 and 1e-16 are investigated.

**Figure 6.**
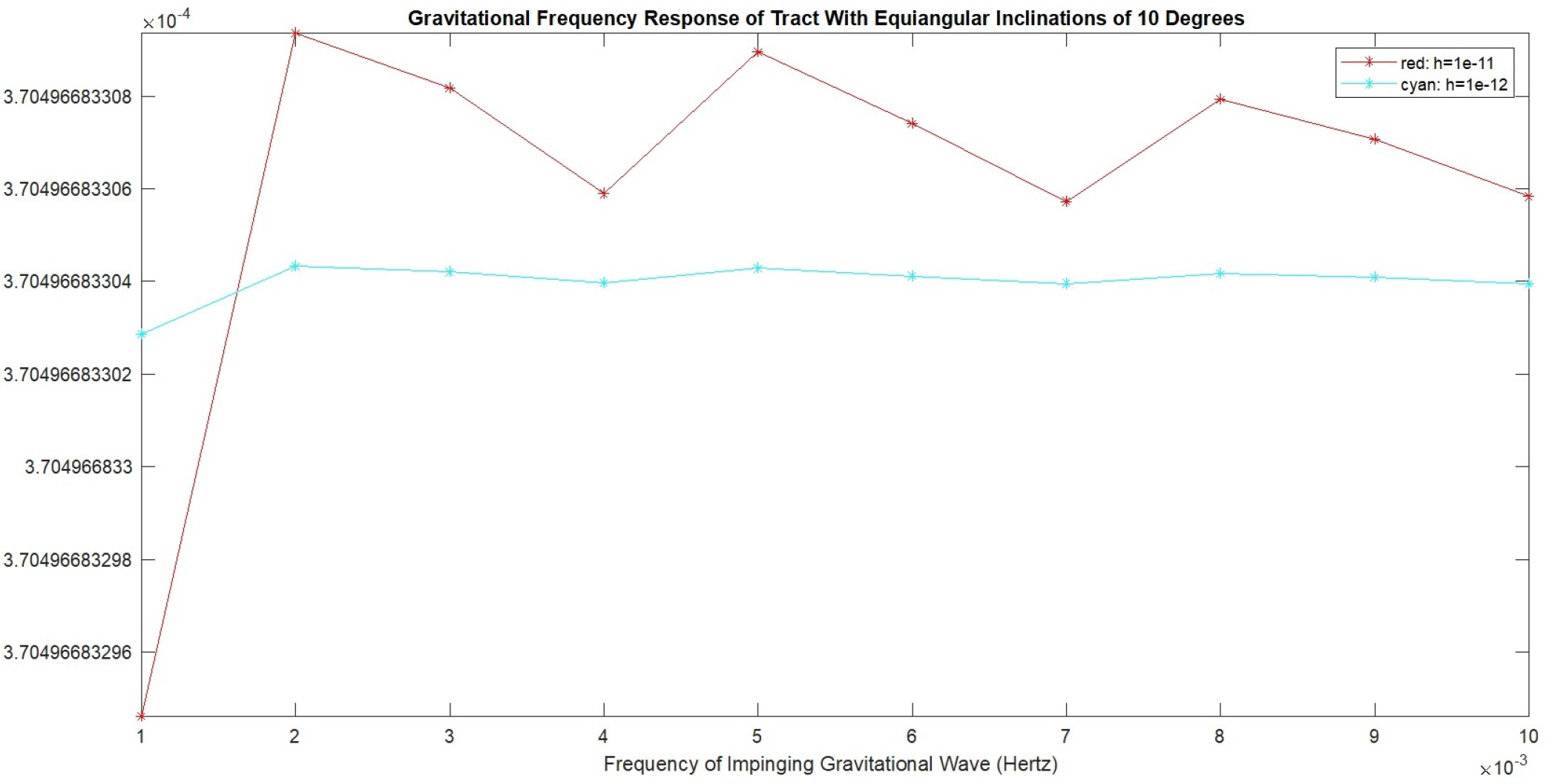
In figure 6, the x-axis is the frequency of the impinging gravitational wave in Hertz, ranging from 1e-3 Hz to 1e-2 Hz. The y-axis is the magnitude of the temporal difference (in seconds) between action potential initiation times on axons 2 and 3 out of a total of 3 axons. Only 1 axon was stimulated, but all 3 are ephaptically coupled within the tract. Each tract considered has a specific geometry. Since the tract was considered in the absence of any ion-channel or other noise source, these are deterministic and not probabilistic simulations. As a result these output graphs are not amenable to statistical significance tests. In each figure, the two applied gravitational wave (dimensionless) strain values are shown in the legend boxes and the point-line graphs are correspondingly color coded. Overall in this set of figures, strains between 1e-8 and 1e-16 are investigated.

**Figure 7.**
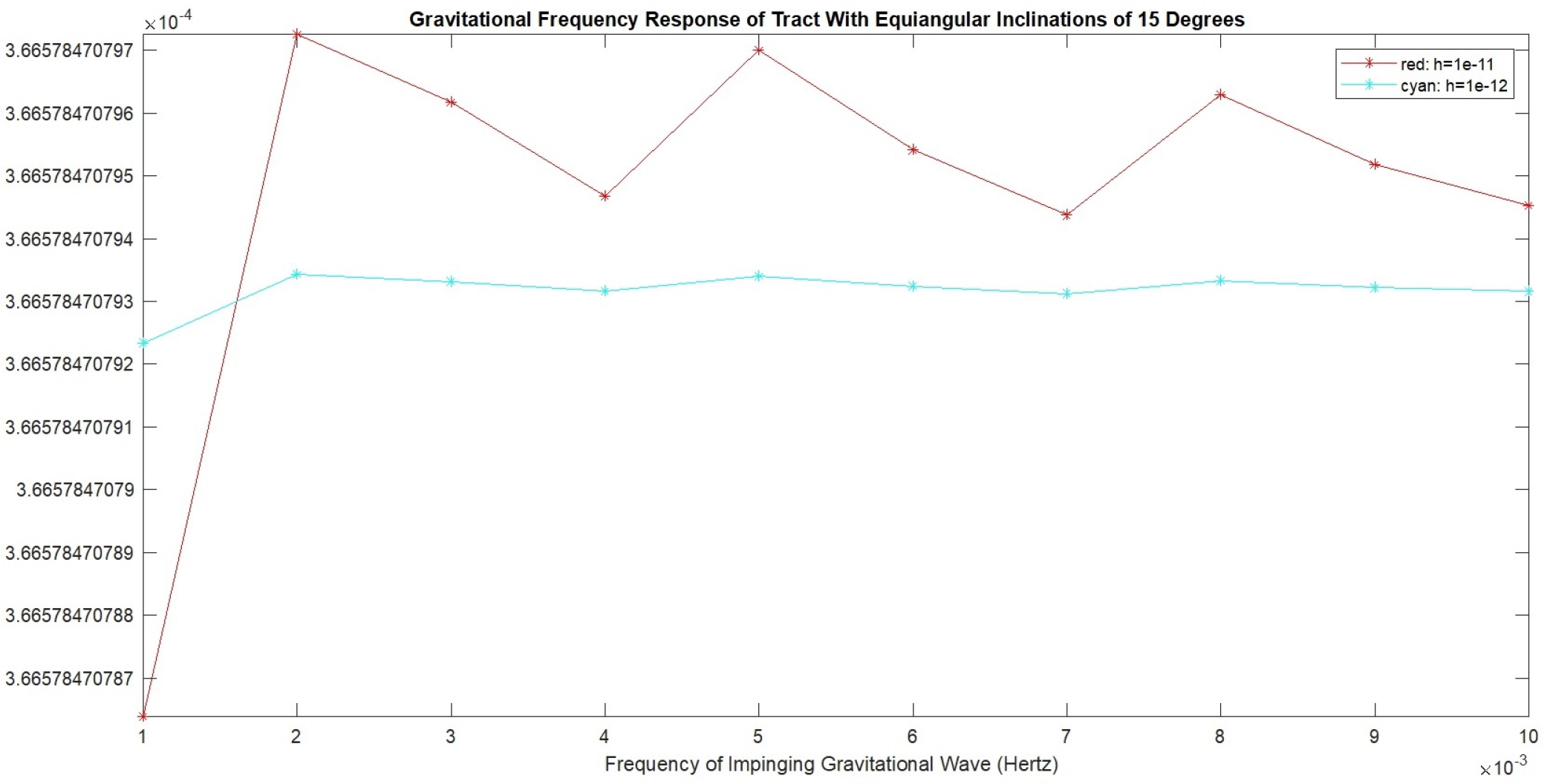
In figure 7, the x-axis is the frequency of the impinging gravitational wave in Hertz, ranging from 1e-3 Hz to 1e-2 Hz. The y-axis is the magnitude of the temporal difference (in seconds) between action potential initiation times on axons 2 and 3 out of a total of 3 axons. Only 1 axon was stimulated, but all 3 are ephaptically coupled within the tract. Each tract considered has a specific geometry. Since the tract was considered in the absence of any ion-channel or other noise source, these are deterministic and not probabilistic simulations. As a result these output graphs are not amenable to statistical significance tests. In each figure, the two applied gravitational wave (dimensionless) strain values are shown in the legend boxes and the point-line graphs are correspondingly color coded. Overall in this set of figures, strains between 1e-8 and 1e-16 are investigated.

**Figure 8.**
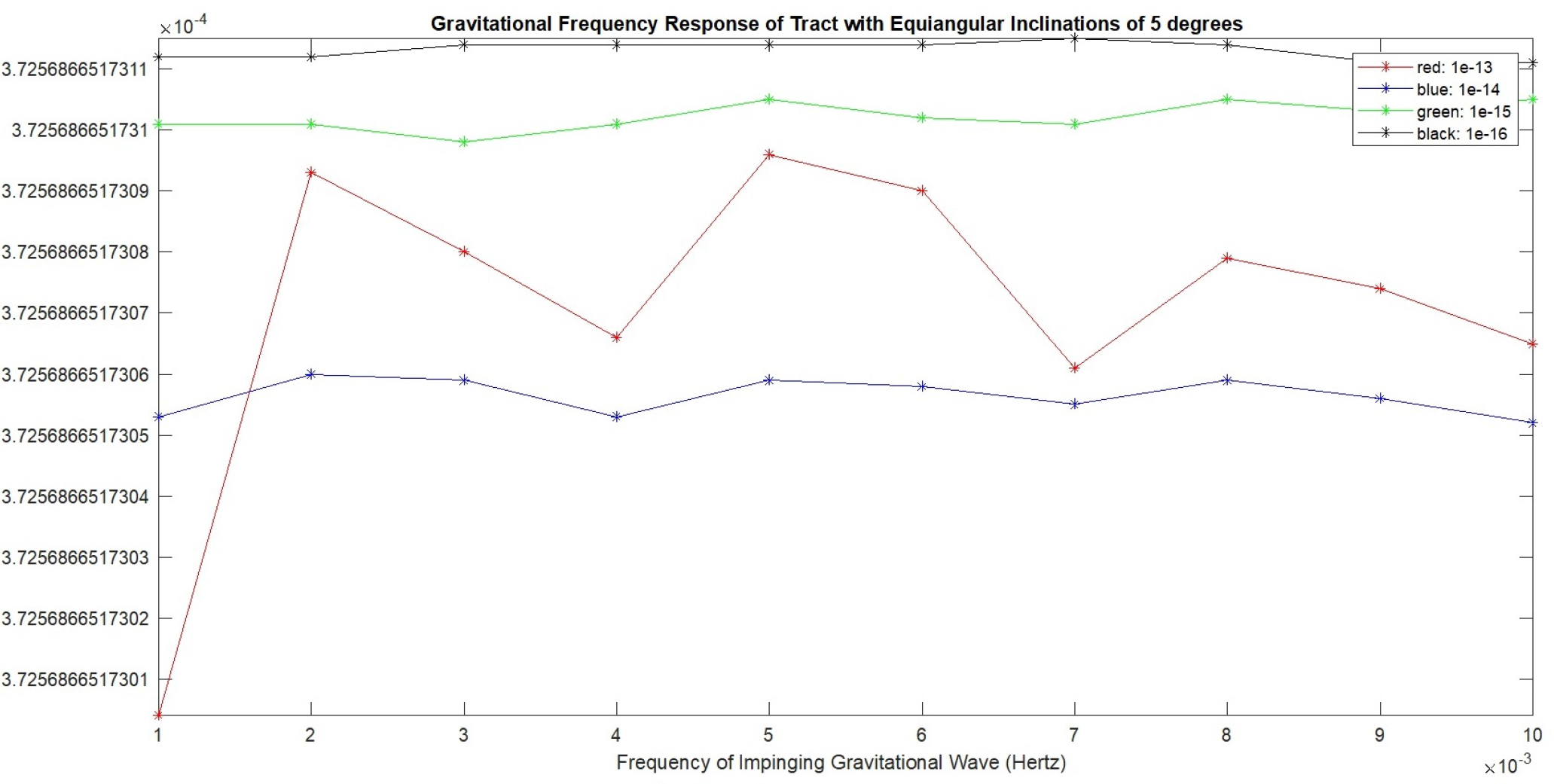
In figure 8, the x-axis is the frequency of the impinging gravitational wave in Hertz, ranging from 1e-3 Hz to 1e-2 Hz. The y-axis is the magnitude of the temporal difference (in seconds) between action potential initiation times on axons 2 and 3 out of a total of 3 axons. Only 1 axon was stimulated, but all 3 are ephaptically coupled within the tract. Each tract considered has a specific geometry. Since the tract was considered in the absence of any ion-channel or other noise source, these are deterministic and not probabilistic simulations. As a result these output graphs are not amenable to statistical significance tests. In each figure, the two applied gravitational wave (dimensionless) strain values are shown in the legend boxes and the point-line graphs are correspondingly color coded. Overall in this set of figures, strains between 1e-8 and 1e-16 are investigated.

**Figure 9.**
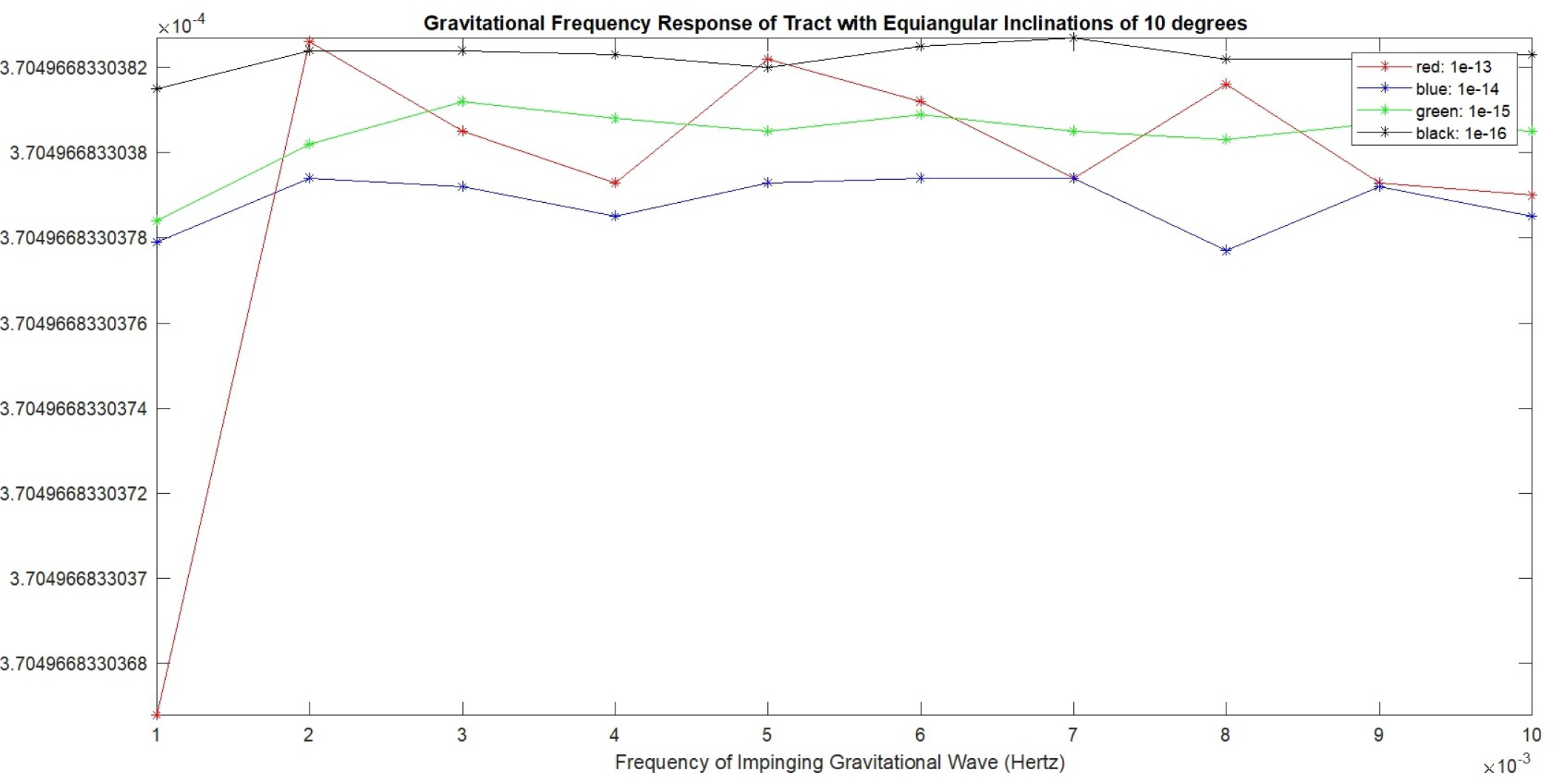
In figure 9, the x-axis is the frequency of the impinging gravitational wave in Hertz, ranging from 1e-3 Hz to 1e-2 Hz. The y-axis is the magnitude of the temporal difference (in seconds) between action potential initiation times on axons 2 and 3 out of a total of 3 axons. Only 1 axon was stimulated, but all 3 are ephaptically coupled within the tract. Each tract considered has a specific geometry. Since the tract was considered in the absence of any ion-channel or other noise source, these are deterministic and not probabilistic simulations. As a result these output graphs are not amenable to statistical significance tests. In each figure, the two applied gravitational wave (dimensionless) strain values are shown in the legend boxes and the point-line graphs are correspondingly color coded. Overall in this set of figures, strains between 1e-8 and 1e-16 are investigated.

**Figure 10.**
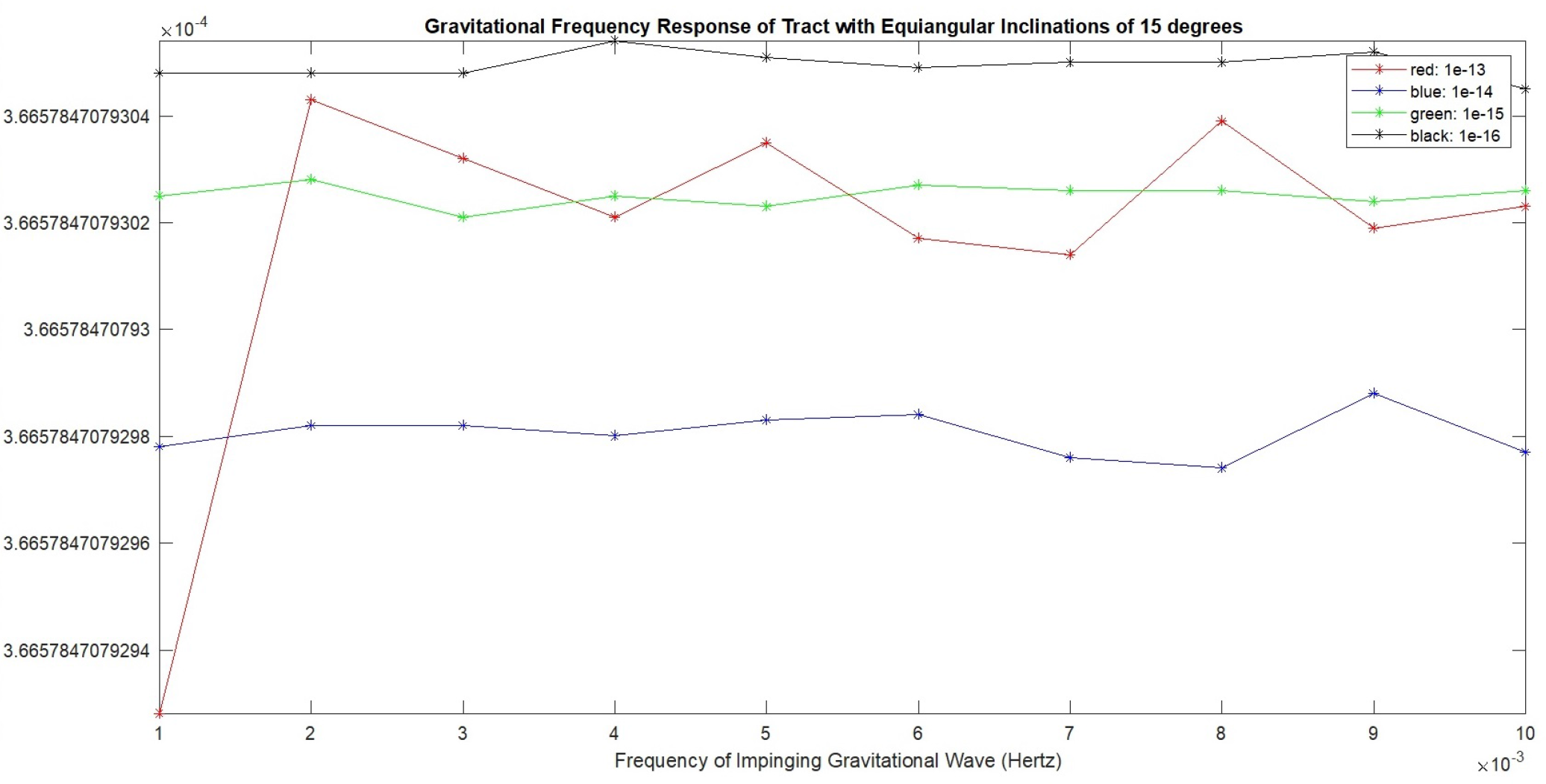
Figure 10 has the exact same x- and y-axis specifications as the other figures. The inclination considered in this figure is 15 degrees. The strain values investigated in this figure are 1e-13, 1e-14, 1e-15 and 1e-16.

## 4 Discussion

A comparative look at Figures 2 through 10 indicates that the response depends on strain, frequency as well as the local geometry of the tract. It is also clear that a strain being lower in magnitude than another, doesn’t guarantee a weaker response. Further it is observed that an angle being larger than another doesn’t guarantee increased discriminability amongst strains. There is thus an intimate (though not direct) relationship between the gravitational wave’s properties and the response of the tract in different configurations. This indicates that measuring this response, say by aid of optogenetics, can give some clue to metric perturbations in the vicinity.

## 5 Conclusion

This result has ramifications for man’s understanding of himself and his place in the Universe. For example, under the materialist philosophical view [1] that the mind is physical (or, more clearly, bio-physical), when action potential timing is impacted to some degree by distant events in spacetime, it constitutes an impact on man’s thoughts and, consequently, man’s behavior.

We leave it to our readers to investigate this brain-cosmos coupling in more detail and flesh out the implications of this finding. As a parting note, we would like to point out the strange and rather unsettlingly close analogy of this result with the Vedic idea that individual soul (*atman*) is identical with cosmic soul (*Brahman*) [4].

